# GAT1508, a selective neuronal GIRK1/2 activator, exhibits superior antiepileptic efficacy and synergizes with cannabidiol

**DOI:** 10.64898/2026.10.14.758970

**Authors:** Jahnavi Simhadri, Andrew Zorn, Liang Wang, Takeharu Kawano, Natalia Cordero-Ruiz, Lida Vagiaki, Nousheen Parvaiz, Mehreen Zaka, Lucas Cantwell, Kim W. Chan, Ana Santa-Cruz, Evangelos Dadiotis, Nicole Rivera, Wilder Felix, Ganesh A. Thakur, Roman Manetsch, Leigh D. Plant, Kyriaki Sidiropoulou, Meng Cui, Georgios Skiniotis, Diomedes E. Logothetis, Stelios M. Smirnakis

## Abstract

G protein-gated inwardly rectifying potassium (GIRK; Kir3) channels are key regulators of neuronal excitability and represent promising therapeutic targets for epilepsy. Here, we characterize GAT1508, a urea-based small-molecule activator selectively targeting neuronal GIRK1/2 heterotetramers, and compare its efficacy with previously described urea compounds ML297 and GiGA1. In a pentylenetetrazole (PTZ)-induced seizure model, GAT1508 demonstrated robust antiseizure activity, outperforming both ML297 and GiGA1 and achieving efficacy comparable to valproic acid. Electrophysiological studies using two-electrode voltage clamp in *Xenopus* oocytes revealed that GAT1508, unlike ML297 and GiGA1, selectively activates GIRK1/2 channels without significant activation of cardiac GIRK1/4 channels, suggesting reduced risk of cardiovascular side effects. Structural and computational analyses indicate that enhanced stabilization of GAT1508 compared to ML297 within the GIRK1/2 binding pocket promotes both ion permeation and channel gating through coordinated interactions with key residues. Furthermore, GAT1508, like the two other urea drugs, exhibited synergistic activation of GIRK channels when combined with Gβγ-mediated pathways and significantly potentiated the antiseizure effects of cannabidiol (CBD) in both hippocampal slice recordings and *in vivo* seizure models. These findings establish GAT1508 as a potent and selective GIRK1/2 activator and support a novel combination therapeutic strategy for epilepsy based on convergent GIRK channel modulation.

---

G protein-gated inwardly rectifying potassium (GIRK; Kir3) channels are critical determinants of cellular excitability, particularly in neurons, where they stabilize the resting membrane potential and dampen excitatory signaling. GIRK channels are activated by Gβγ subunits released from Gi/o-coupled G protein-coupled receptors (GPCRs) (1, 2), intracellular Na^+^ (3), and require the membrane phospholipid phosphatidylinositol 4,5-bisphosphate (PIP_2_) as an essential cofactor (4, 5). Although PIP_2_ alone does not efficiently open GIRK channels, it is required for activation by gating molecules including Gβγ, intracellular Na^+^, and urea-based activators such as ML297, GAT1508, and GiGA1 (6, 7). Functional GIRK channels form as heterotetramers, typically comprising GIRK1 with GIRK2 or GIRK4 subunits, with GIRK1/2 channels predominating in the central nervous system (CNS) (8).

By maintaining membrane potential near the potassium equilibrium potential, GIRK channels suppress neuronal firing by limiting activation of voltage-gated Na^+^ and Ca^2+^ channels. Dysregulation of GIRK activity has been implicated in multiple neurological disorders. Reduced GIRK function is associated with neuronal hyperexcitability and epilepsy, whereas excessive activity has been linked to cognitive deficits such as those observed in Down syndrome (9, 10).

Functional GIRK channels contain multiple constriction sites along the ion-permeation pathway. The selectivity filter (SF) confers K^+^ selectivity, whereas the helix-bundle crossing (HBC) and G-loop form two regulated gates in series. The HBC gate is formed near the inner leaflet of the plasma membrane, and the cytosolic G-loop gate is characteristic of Kir channels. Both gates must open to support ion conduction (6).

Molecular dynamics (MD) simulations performed in the presence of an applied electric field have shown that regulators acting predominantly at distinct channel-gating sites can act synergistically. In these simulations, the GIRK2–Gβγ–Na^+^ system produced greater gate opening and ion permeation than either GIRK2–Gβγ, which predominantly regulates the HBC gate, or GIRK2– Na^+^, which regulates the G-loop gate, alone (6). These findings were consistent with earlier experimental results (11). Moreover, a small-molecule urea activator was shown to synergize with GABA_B_ receptor stimulation to activate GIRK currents in basolateral amygdala slices (12).

Small-molecule GIRK activators represent a promising therapeutic strategy. ML297, the first selective GIRK1 heterotetrameric channel activator demonstrating anticonvulsant properties in preclinical models (13), was followed by GiGA1 (14). However, both compounds exhibit limited subtype selectivity, activating cardiac GIRK1/4 channels and raising safety concerns. GAT1508, a structural analog of ML297, was developed to preferentially target neuronal GIRK1/2 channels (12).

In the present study, we demonstrate that GAT1508 exhibits superior antiseizure efficacy compared with ML297 and GiGA1, elucidate its mechanism of action, and show that it synergizes with GPCR-mediated GIRK activation. Notably, we identify a functional synergy between GAT1508 and cannabidiol (CBD), an FDA-approved antiseizure agent, supporting a novel combinatorial therapeutic strategy for epilepsy.

## Results

### GAT1508 exhibits superior antiepileptic efficacy over reported GIRK activators and other antiepileptics

Urea-based GIRK activators, ML297 and GiGA1 have previously demonstrated efficacy in preventing seizures in acute seizure models in mice (13, 14). GAT1508, is a bromo-thiophene substituted small molecule derived from ML297, which was found to be highly selective for the neuronal GIRK1/2 channels without significantly affecting GIRK1/4 channels predominantly expressed in heart minimizing cardiac side effects (12).

Our first aim was to compare the antiseizure efficacy of GAT1508 with the existing urea compounds ML297 and GiGA1. To this end, we conducted pilot studies in wild-type (WT) C57BL/6 mice from which we determined that GAT1508 showed robust antiseizure properties at 60 mg/kg. Previous studies reported similar effective doses for ML297 and GiGA1 (13, 14). All three drugs were compared in an acute seizure model, using a chemoconvulsant, pentylenetetrazole (PTZ, 60 mg/kg, i.p., 10 mg/ml). C57BL/6 mice were pretreated with previously effective doses of 60 mg/kg of GAT1508, GiGA1, ML297 (12-14) or vehicle control 30 min prior to PTZ administration (Fig. 1*A*). Video recordings of the mice were taken for 30 min following PTZ administration and seizure severity was scored using a modified Racine score (Table S1), with grades greater than or equal to four (score 4) representing tonic-clonic seizures.

**Figure 1.**
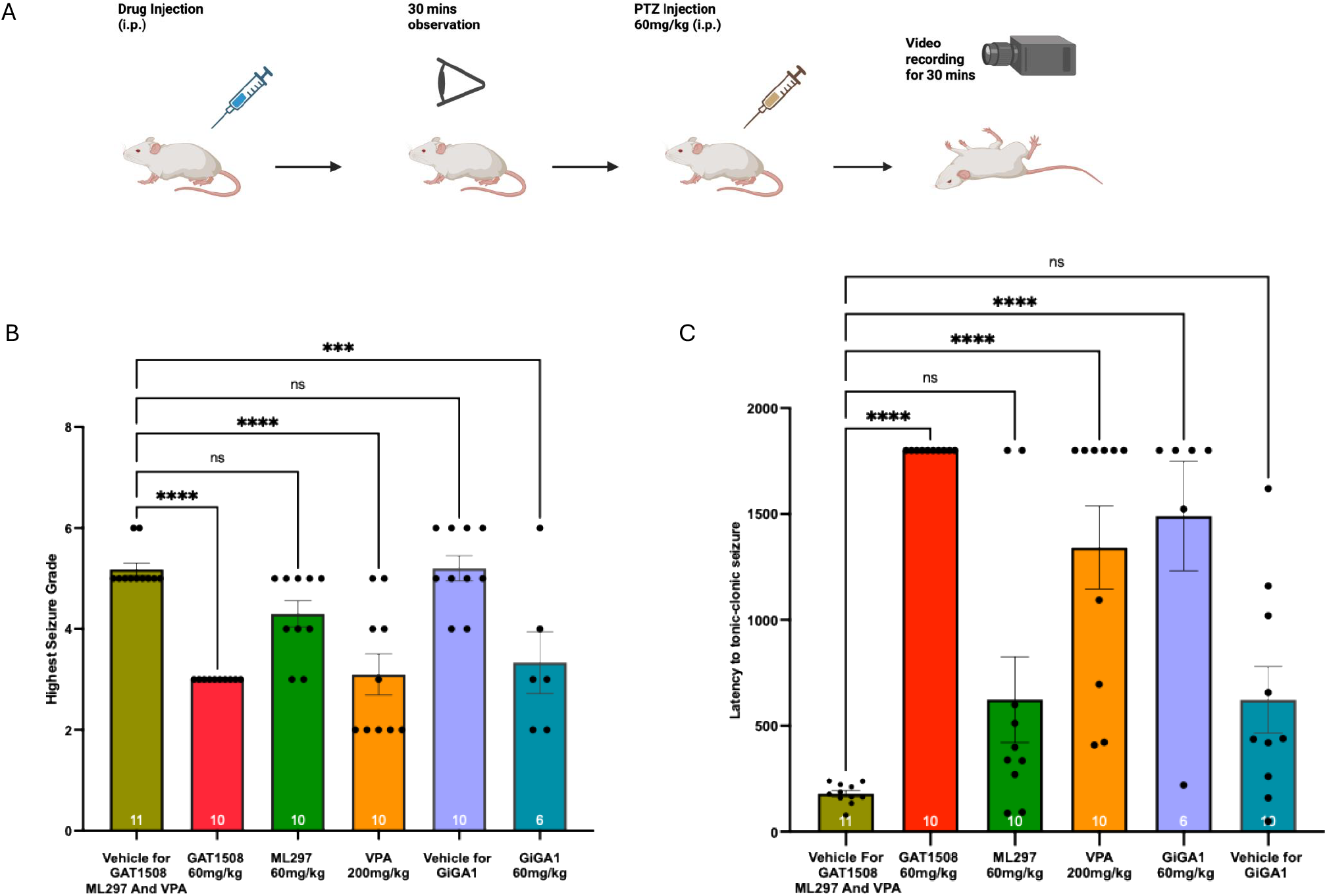
GAT1508 exhibits superior antiseizure properties compared to other urea-based GIRK channel activators. *A*, WT C57BL/6 mice (Charles River Laboratories) were injected intraperitoneally (i.p.) with either of the drugs GAT1508, ML297, GiGA1 at 60 mg/kg followed by a pro-convulsant dose of PTZ 60 mg/kg i.p. *B*, Shows the highest grade of seizure observed in each treatment group and *C*, Latency (in seconds) to the tonic-clonic seizures for each cohort. Statistical significance was tested using ordinary one-way ANOVA where P***<0.001 and P****<0.0001.

GAT1508 treated mice showed no seizure above grade 3 (Fig. 1*B, C*). The results from GAT1508 60 mg/kg were comparable, albeit less variable, to valproic acid (VPA) 200 mg/kg, the gold-standard antiseizure drug which served as a positive control. In our hands, ML297 did not show efficacy as an antiseizure agent as it was not significantly different from the vehicle in preventing seizures and neither was it able to delay seizures significantly compared to vehicle controls. Due to solubility differences GiGA1 was delivered in a different vehicle than the other urea compounds. Although GiGA1 showed some significant difference from its vehicle, the results were highly variable as mice treated with GiGA1at 60 mg/kg exhibited sedation and respiratory depression. These complications limited our ability to accurately grade the seizures in this treatment group, as several mice died without any apparent tonic-clonic seizures. While death was scored as Grade 6 on the Racine scale, the absence of an apparent tonic-clonic seizure precluded reliable evaluation. Thus, even though double the number of mice (n=12) were tested in this cohort half the results are not included due to death (n=6).

To address this limitation, we compared GAT1508 and GiGA1 at a reduced dose of 30 mg/kg (Fig. S1). Even at lowered doses, some mice receiving GiGA1 showed signs of sedation, and once again the results from GiGA1 were quite variable, unlike GAT1508 30 mg/kg that consistently showed protection from seizures above grade 4. Taken together, these results suggest that GAT1508 is comparable to valproate and the most effective anti-seizure compound amongst the three urea-based GIRK activators tested in our study.

### Specificity of the Urea compounds

We next assessed the subtype selectivity of the urea compounds. Selective activation of the neuronal GIRK1/2 channels over cardiac GIRK1/4 is critical to avoid/prevent cardiac side effects such as bradycardia, hypotension and cardiac arrythmias seen with GIRK1/4 activation. Using two-electrode voltage clamp (TEVC), we compared activity of GAT1508, ML297 and GiGA1 in oocytes expressing either GIRK1/2 or GIRK1/4 subtype channels.

GiGA1 was found to be the least selective, showing comparable activation of both GIRK1/2 as well as GIRK1/4 channels (Fig. 2*G*–*I*). ML297 exhibited modest preference for GIRK1/2 channel but still showed a noticeable degree of activation of GIRK1/4 (12, 15; Fig. 2*D*–*F*). GAT1508, selectively targeted GIRK1/2 (12) with no significant effect on GIRK1/4 channels (I/I_basal_, i.e, no significantly induced current for GIRK1/4 channel subtype) (Fig. 2*A*–*C*).

**Figure 2.**
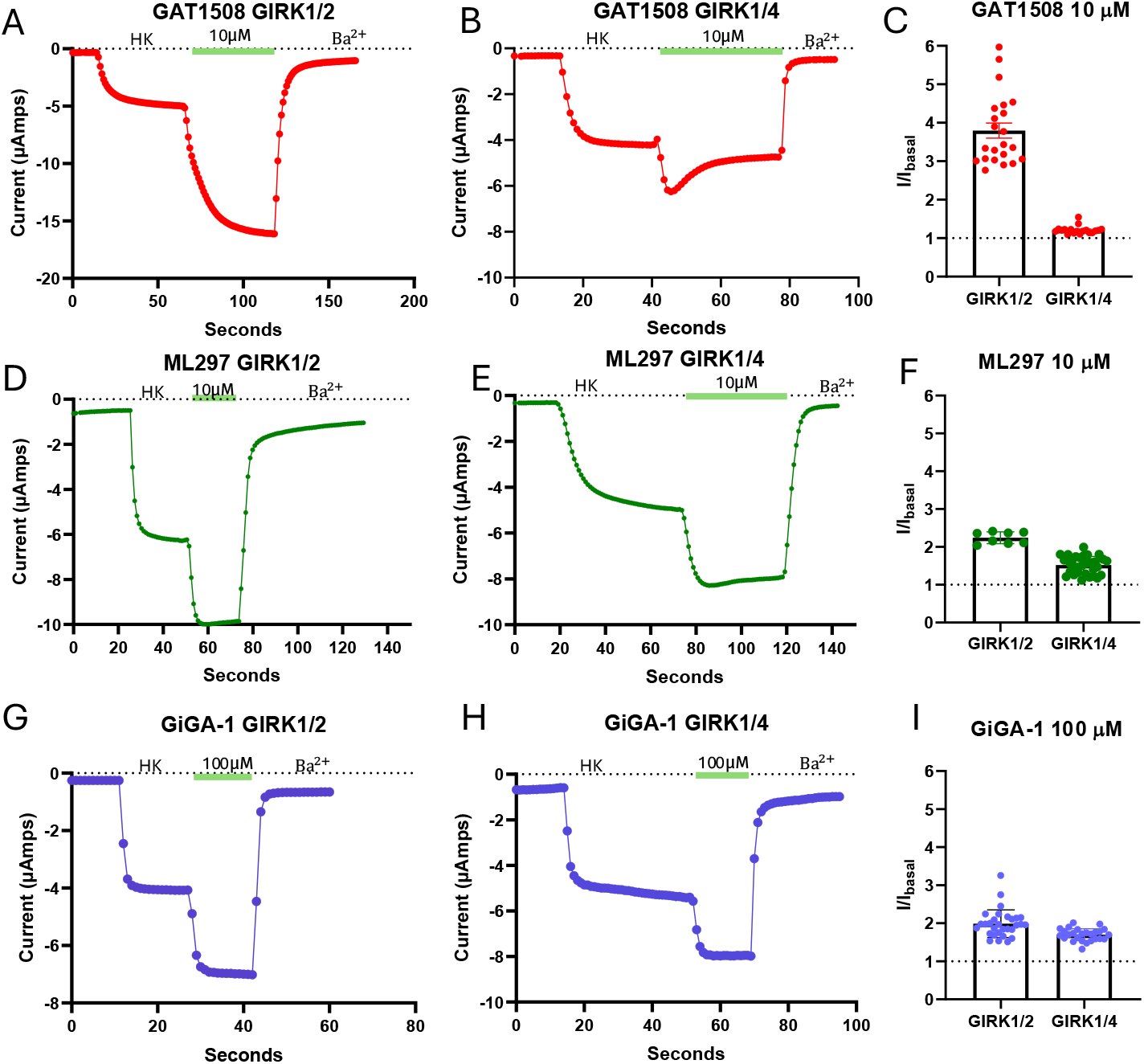
GAT1508 selectively activates neuronal GIRK1/2 channels, unlike ML297 and GiGA1. Two-electrode voltage-clamp recordings from Xenopus oocytes were used to assess subtype selectivity. *A–C*, 10 μM GAT1508 robustly activates GIRK1/2 (n = 22; EC_50_ ≈ 75 nM) with minimal effect on GIRK1/4 (n = 18). *D–F*, 10 μM ML297 activates both GIRK1/2 (n = 9; EC_50_ ≈ 110 nM) and GIRK1/4 (n = 32; EC_50_ ≈ 1.2 μM) above basal current. *G–I*, 100 μM GiGA1 (EC_50_ ≈ 30 μM) activates GIRK1/2 (n = 30) and GIRK1/4 (n = 26) comparably. The dotted line indicates basal current. Concentrations shown are saturating concentrations derived from the corresponding dose-response curves.

### The GAT1508 binding site and the importance of each interaction

We have reported a high-resolution atomic structure of ML297 in the GIRK1/2 binding pocket (16). Here, we report the complex of this asymmetric structure composed of 1GIRK1:3GIRK2 subunits bound to GAT1508 (Fig. 3*A*). The GAT1508 binding site is comprised by four contacts of the drug with the GIRK1 subunit (the oxygen of the urea carbonyl with W95.1, the hydrogen of the urea amides interact dynamically with either the D173.1 or the E141.1, and the tail bromine with M98.1 or the sulphur of the thiophene with the imidazole of W95.1). One contact is made with the GIRK2 subunit (the nitrogen of the pyrazole with S177.2) (Fig. 3*B*). The workflow for the cryo-EM structure determination of GIRK1/2 in complex with GAT1508 at 2.7 Å resolution is shown in Fig. S2.

**Figure 3.**
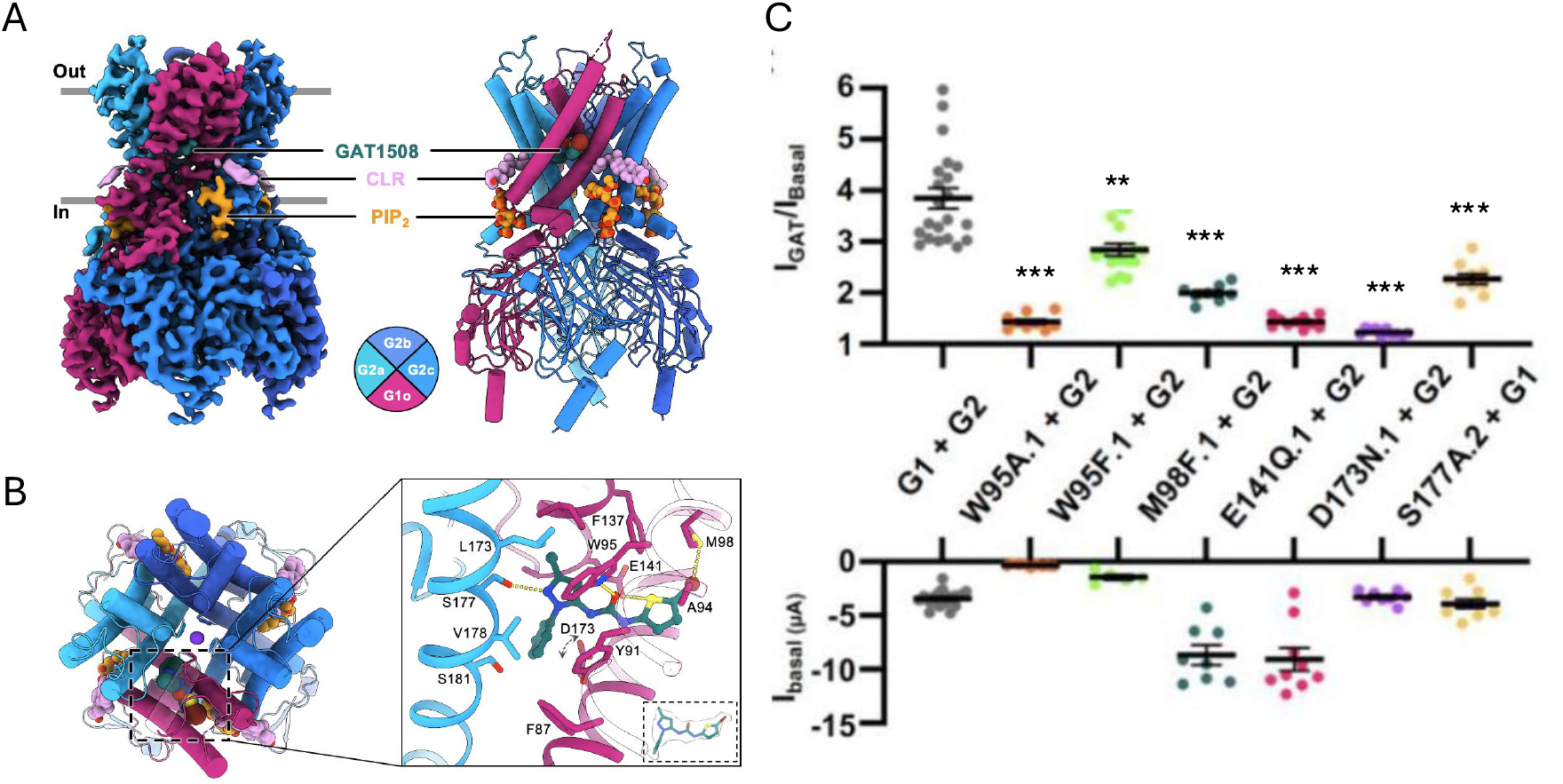
GAT1508 binds a GIRK1-specific transmembrane pocket to stimulate ion flow. *A*, Cryo-EM side view structure of the asymmetric GIRK1/2, with one GIRK1 (red) and 3 GIRK2 (blue) bound to the activator GAT1508, Cholesterol (CLR) and di-eight carbon long acyl chain phosphatidylinostol bis-phoshate (diC8-PIP_2_) (top). *B*, The GAT1508 binding site seen from the extracellular side (left) with a zoomed in close up of the interactions of the ligand in its binding pocket. The ligand occupies a pocket formed primarily by the single GIRK1 subunit (G1s; red) with contributions from the adjacent G2a (a,b,c clockwise; blue). Double-headed arrows indicate the conformational mobility of the Asp side chain (D173). *C*, Summary current of activation in response to 10 μM GAT1508 (I_GAT_) over basal current (I_Basal)_ represented as I_GAT_/ I_Basal_ in *Xenopus* oocytes expressing GIRK1 + GIRK2 (G1 + G2) with the mutations indicated, in the GIRK1 or the GIRK2 subunit. Basal currents are inward (negative) currents at High K^+^ symmetrical solutions. Statistical significance is assessed compared to the wild type (G1 + G2) subunits. [[describe the bottom row – basal current ; also the mutations briefly or refer to the text]] Statistical significance was assessed using a two-tailed unpaired Welch’s t test where **p < 0.01 and ***p < 0.001.

D173.1 interactions with the drug were dynamic and could not be captured by the cryo-EM structure. Mutagenesis of all five binding sites (W95F.1, M98F.1, E141Q.1, D173N.1 and S177A.2) confirmed their importance as the ability of GAT1508 to induce currents over basal activity (i.e., in the absence of ligand) was significantly reduced (Fig. 3C).

The higher potency of GAT1508 than ML297 (12) was corroborated by Free Energy Perturbation (FEP) calculations of 10 urea and acetamide ligands using the FEP+ module of the Schrödinger software (Schrödinger Release 2024-4) following a standard protocol (17). In this assessment of ligand energy of interaction in the binding pocket the pKi of GAT1508 was estimated to be 7.8 ± 0.32 compared to ML297 of 6.5 ± 0.36 (Fig. 4).

**Figure 4.**
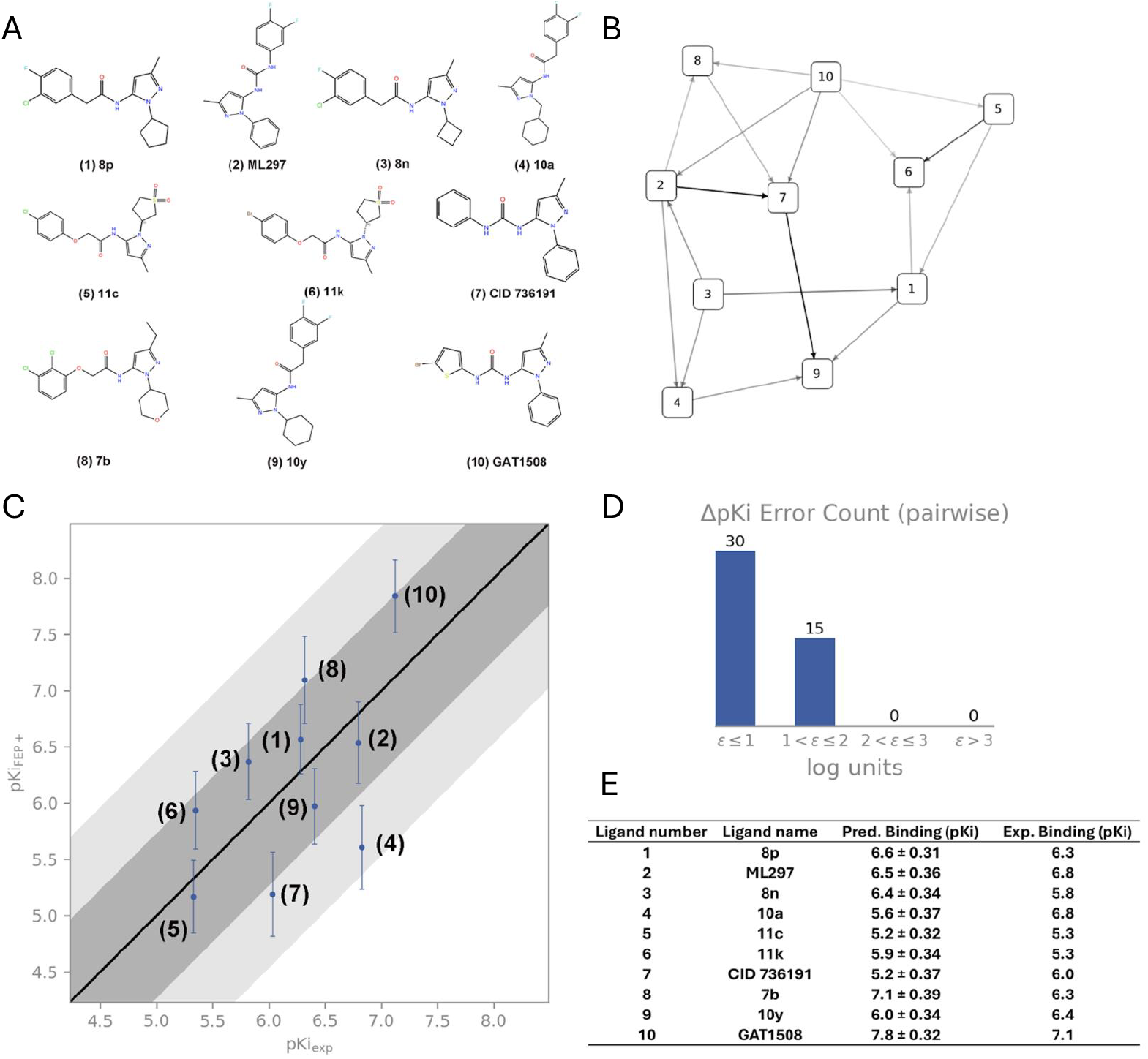
Relative binding free energies between the drug molecules and the GIRK1/2 channel predicted by FEP (Free Energy Perturbation) calculations. *A*, 2D structures of ML297 and selected analogs. *B*, Cycle-closure FEP map, where each node represents a compound and each arrow indicates an alchemical transformation between two molecules. *C*, Correlation plot comparing experimental (X-axis) and FEP-predicted (Y-axis) binding affinities for the GIRK1/2 channel. MUE (pairwise mean unsigned error) = 1.12; RMSE (pairwise root mean squared error) = 1.24; Expected Exp. R^2^ = 0.57; Expected Pred. R^2^ = 0.32; Predicted R^2^ = 0.34. *D*, ΔΔpKi error distribution for the FEP results: bin 1, ΔΔpKi ≤ 1(excellent prediction); bin 2, 1 < ΔΔpKi ≤ 2 (acceptable prediction); bin 3, 2 < ΔΔpKi ≤ 3 (poor prediction); bin 4, ΔΔpKi > 3 (failed prediction). E, Data table corresponding to panel *C*.

The stabilization of GAT1508 in the binding pocket, as suggested by the FEP calculations, resulted in changes in the frequency of both stable and dynamic interactions of the ligand with key residues in the pocket. Gating molecular dynamics (GMD) simulations are typical MD simulations that are able to capture the opening of the channel gates and ion permeation over the course of the simulation run (18). This allows quantification of the time specific interactions occur over the course of the simulation and the percentage this interaction occurs over the total simulation time (ST). Additionally, quantification is also possible of the number and percentage (over all ions permeating during the run) of ions permeating (PC: permeation count) while a particular key interaction occurred.

A GMD comparison of GAT1508 with ML297 bound in the pocket as well as with the unliganded GIRK1/2 channel (GIRK1/2 Apo) reveal that the bromothiophene stabilization increased dramatically the interaction of the ligand with W95.1 and prevented flipping of the ligand so that the amides of the urea scaffold interacted with D173.1 rather than the E141.1 and the nitrogen of the pyrazole with S177.2 rather than with S181.2 (Fig. 5). These interaction changes resulted in the following sequence of number of ions permeating (highest to lowest): GAT1508 > ML297 > GIRK1/2 Apo.

**Figure 5.**
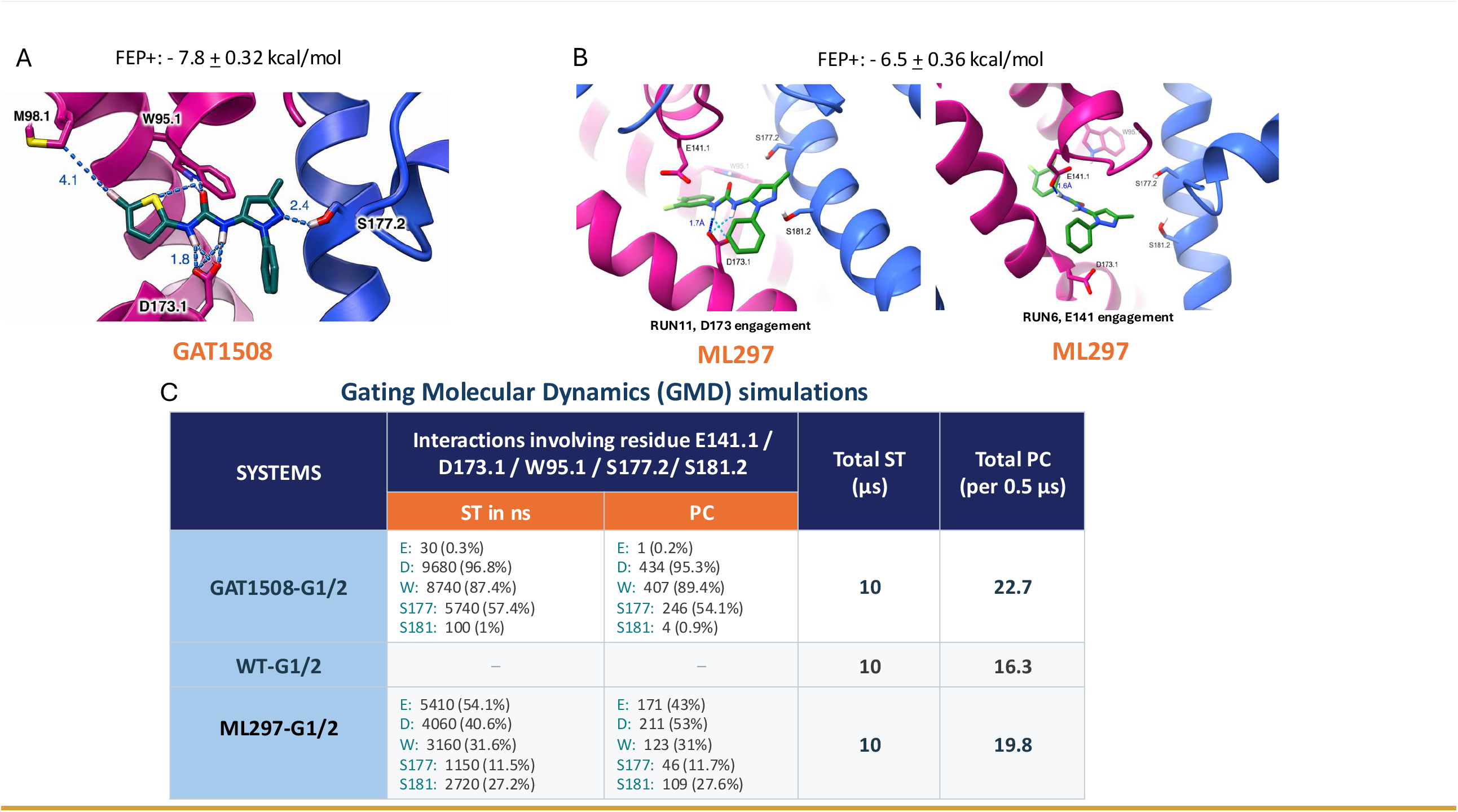
GMD simulations provide insight on the dynamic behavior of D173.1 and the interactions of the four remaining key residues *A:* A representative conformation of GAT1508 highlighting the dominant pattern of hydrogen-bonding geometry in the GIRK1/2 binding pocket displaying the stable interaction of the ligand in the binding pocket. GAT1508 binds stably in the GIRK1/2 pocket stabilizing the D173.1 and S177.2 interactions. This is accomplished by the bromothiophene tail stabilizing GAT1508 in the binding pocket through interaction of the bromine with M98.1, while the sulphur of the thiophene interacts with the imidazole of W95.1. This stabilization of the ligand in the pocket enhances all its interactions (with W95.1, D173.1 and S177.2). *B*, ML297 is not as stable in the binding pocket, flipping to enable interaction of its amides in the urea scaffold with either D173.1 or E141.1 (also interactions of the nitrogen of its pyrazole with either S177.2 or S181.2 - not shown). *C*, Summary table showing the Simulation Time (ST) of the Apo (unliganded) GIRK1/2 (WT-G1/2) as well as specific residue interactions of GIRK1/2 with GAT1508 (GAT-G1/2) that were encountered in the course of several 500 ns-long runs amounting to an overall simulation time of 10 µs for WT-G1/2 and 10 µs for GAT-G1/2. Corresponding permeation counts (PC) and percentages are also shown. A representative conformation of GAT1508 highlighting the dominant pattern of hydrogen-bonding geometry in the GIRK1/2 binding pocket displaying the stable interaction of the ligand in the binding pocket.

### GAT1508 synergizes with GPCR agonists to activate the GIRK1/2 channel

Building on gating synergies reported previously (6, 11, 12), we investigated if a similar synergistic interaction would occur in in vitro, ex vivo and in vivo systems between the urea-based GIRK activators like GAT1508 and compounds acting on distinct binding sites, such as GPCR agonists that activate the GIRK channel via Gβγ. We first tested this synergy between the urea compounds and 1-Propanol, which shares a similar binding site as Gβγ.

Since the synergy primarily manifests itself in potency changes rather than the maximal efficacy, we performed dose-response curve experiments to test how each compound affects the potency of 1-Propanol. Dose-response curves of 1-Propanol were generated in the absence and presence of the minimum effective concentrations (MEC; Fig. S3) and the EC_50_ values were compared (Fig. 6). In the GAT1508 experiment, the EC_50_ of 1-Propanol alone was 81 mM, whereas co-application of 100 nM GAT1508 shifted it to 54 mM. Similarly, in the ML297 experiment the EC_50_ of 1-Propanol alone shifted from 68 mM to 39 mM when co-applied with 100 nM ML297. A comparable leftward shift was also observed with GiGA1, where the EC_50_ of 1-Propanol decreased from 84 mM to 45 mM. In contrast baclofen, taken as a negative control since it acts at the same binding site as 1-Propanol did not significantly alter the EC_50_ (1-Propanol: 84 mM; 1-Propanol + 100 nM Baclofen: 86 mM) failed to exhibit synergy.

**Figure 6.**
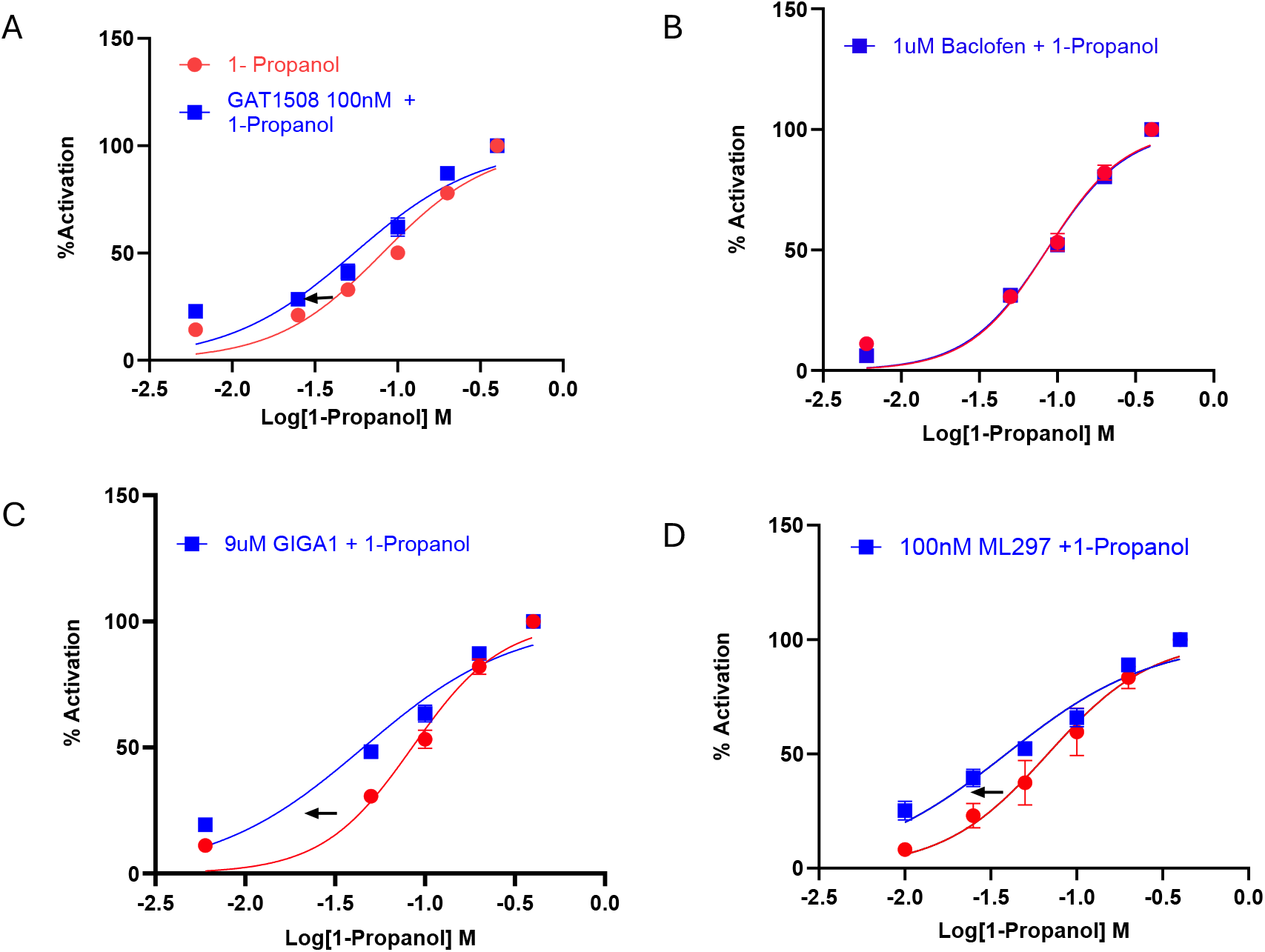
Urea-based activators synergize with 1-Propanol, likely by acting on a distinct binding site, whereas Gβγ (via baclofen) does not. A minimal effective concentration (MECs) (taken from individual dose-response curves; see Fig. S2) were co-applied with increasing doses of 1-Propanol. Resulting dose-response curves of 1-Propanol co-applied with compounds were compared with the dose-response curve of 1-Propanol alone. Urea compounds ML297, GAT1508 and GiGA1 co-application with 1-Propanol produced a leftward shift in the 1-propanol dose-response curve reducing the EC_50_ of 1-Propanol *A*, with GAT1508 *C*, with GiGA1 and *D*, with ML297. In contrast, co-application of *B*, baclofen with 1-Propanol did not exhibit synergism.

### Countering seizure activity in ex vivo and in vivo models of epilepsy through synergism

We then explored whether this mechanistic insight of synergism could be translated to a pathophysiological context relevant to epilepsy. CBD is an FDA-approved adjunctive treatment for refractory seizures associated with Dravet Syndrome, Lennox-Gastaut Syndrome, and Tuberous Sclerosis Complex. Although the precise mechanism by which CBD acts remains incompletely understood, it is quite possible that CBD acts via a GPCR to activate GIRK channels and promote neuronal hyperpolarization.Seizure-like events (SLEs) were evoked in hippocampal brain slices using high-K^+^ aCSF (7.5 mM), allowing us to study the intact neuronal networks in the CNS. Field excitatory post-synaptic potentials (fEPSPs) were recorded in normal aCSF, high-K^+^ aCSF and drug diluted in high-K^+^ aCSF. The percent decrease in spontaneous activity in the brain slices with the two drugs (GAT1508 and CBD) was recorded. Fig. 7*A* shows (fEPSP) produced in the presence of high-K^+^ and the effects of high-K^+^ + GAT1508, high-K^+^ + CBD and high-K^+^ + GAT1508 + CBD (5 μM each) on the high-K^+^ induced SLE activity. As indicated in Fig.7 *B, C*, the co-perfusion produced 96.8% inhibition, substantially exceeding both the Bliss prediction (63.1%) and the additive sum of inhibition of each compound applied individually (78.4%). The interaction term was +33.7 percentage points (BCa 95% CI 26–43; percentile CI 26– 43), with every bootstrap replicate above zero (two-sided p**** < 0.0001), indicating synergy. The co-perfusion left only 3.2% of High K^+^ induced activity remaining, versus the 36.9% predicted under independence. In particular, this corresponds to a greater than 11-fold suppression of residual activity relative to what independent effect would predict. Notably, the concentration of the single drug administration was 10 µM whereas the combination in the co-perfusion used only 5 µM of each, so the combination surpassed the independence prediction despite using half the concentration of each drug.

**Figure 7:**
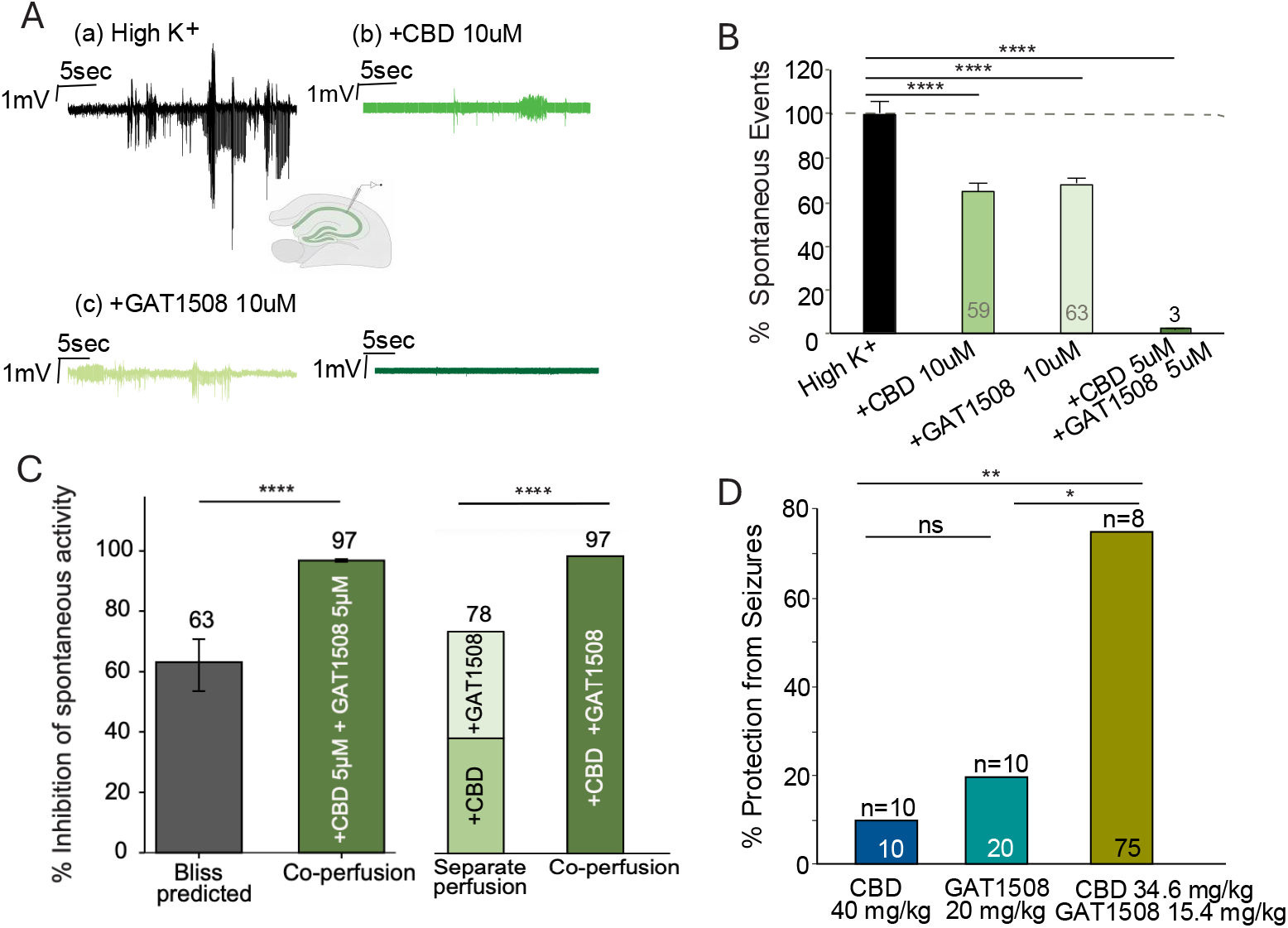
Synergy between CBD and GAT1508 in models of hippocampal slice and in vivo mouse epileptic activity. *A*, fEPSP traces in presence of high-K^+^ (in vitro model of SLE) and the effects of CBD, GAT1508 and CBD + GAT1508 on the SLE activity. *B*, Bar graph showing percentage of spontaneous events and *C*, showing % reduction in the spontaneous activity when CBD or GAT1508 is given alone or in combination showing significant changes from control by co-perfusion to each of the drugs administered alone or in combination. Co-perfusion also shows significant synergy compared to the additive effects of the perfusion of each drug alone. *D*, shows % protection and compares changes for significance from seizures in mice receiving CBD 30 mg/kg, GAT1508 20 mg/kg and CBD + GAT1508 followed by an i.p. injection of a chemoconvulsant PTZ 60 mg/kg.

In vivo experiments on C57BL/6 mice induced with seizures using PTZ also tested synergy between CBD and GAT1508 (Fig. 7*D*). The cohort of mice receiving CBD 30 mg/kg shows no protection from seizures and in the cohort receiving GAT1508 20 mg/kg, 20% of mice showed protection from seizures. Whereas when these two drugs were combined at 34.5 mg/kg of CBD and 15.39 mg/kg of GAT1508, 75% of mice showed significant protection from seizures.

## Discussion

This study establishes GAT1508 as a highly effective and selective activator of neuronal GIRK1/2 channels with significant therapeutic potential for epilepsy. Compared with previously reported urea-based activators, GAT1508 demonstrated superior antiseizure efficacy in vivo, robust subtype selectivity, and a mechanistically distinct mode of action that enhances the most channel gating and ion permeation.

A key finding is the marked functional selectivity of GAT1508 for GIRK1/2 over GIRK1/4 channels. This distinction is critical, as activation of cardiac GIRK1/4 channels can lead to adverse cardiovascular effects, including bradycardia and arrhythmias. Electrophysiological data show that, unlike ML297 and GiGA1, GAT1508 exhibits negligible activation of GIRK1/4 channels, suggesting an improved safety profile for clinical development.

Structural and computational analyses provide mechanistic insight into the enhanced efficacy of GAT1508. Stabilization of the ligand within the GIRK1/2 binding pocket promotes sustained interactions with residues such as D173.1 and S177.2, which appear to coordinate effects on ion permeation and gating. D173.1 and E141.1 are two key residues responsible for the enhanced activity conferred by the GIRK1 subunit (16). Previous reports have suggested that the corresponding residues in GIRK2 homotetramers (N184.2 and E152.2) are determinants of permeation (19, 20). The GIRK1 acidic residues work together to enhance K+ permeation in the heterotetramer compared to the homotetramer GIRK2 (16). S177.2, on the other hand, is one helical turn above the glycine hinge, G180.2, while S181.2 is one residue below. It is likely that interactions of ligands with S177.2 promote TM2 helix flexibility and gating, while interactions with S181.2 constrain flexibility and retard gating. These assertions will need to be further investigated in future studies. Yet, it is clear that with the more stable GAT1508 in the binding pocket, both the S177A.2 and E141Q.1 induced currents were significantly decreased compared with the ML297-induced currents of the same two mutants (Fig. 3*C* and Ref. 16). Importantly, our results demonstrate that GIRK channel activation can be potentiated through synergistic interactions between ligands acting at distinct sites. Urea-based activators, which primarily influence the transmembrane gating machinery, enhance the potency of agents acting through Gβγ-dependent pathways. This is consistent with previous molecular dynamics studies showing that coordinated activation of multiple gating mechanisms produces maximal channel opening (9).

This principle of mechanistic synergy extends to therapeutically relevant combinations. We show that GAT1508 synergizes with cannabidiol (CBD) in vitro, ex vivo and in vivo, significantly enhancing seizure protection at reduced doses. CBD for example activates 5-HT1ARs (21), which have been shown to couple to GIRK channels (22). Given that CBD is limited by dose-dependent adverse effects, including gastrointestinal disturbances and tolerance, combination strategies that reduce required dosing are of considerable clinical interest (23). Our findings suggest that convergent modulation of GIRK channels via distinct molecular pathways represents a viable strategy to improve therapeutic efficacy while minimizing side effects.

In summary, GAT1508 combines high efficacy, subtype selectivity, and synergistic compatibility with existing therapies, positioning it as a promising lead compound for the development of next-generation antiseizure treatments. Future studies will be required to further evaluate its pharmacokinetic properties, long-term safety, and efficacy in chronic epilepsy models.

### Experimental procedures

#### Behavioral studies

PTZ-Induced Acute seizure Paradigm - C57BL/6 male mice were procured from Charles River laboratories; the mice were housed in a temperature and humidity controlled 12-h light-dark cycle room. All experiments performed were approved by the IACUC of the Brigham and Women’s Hospital (BWH). A fresh PTZ stock solution was prepared before each experiment by dissolving the measured amount in 0.9% normal saline at 10 mg/ml concentration. Mice were weighed prior to the experiment, and the dose of the drug and PTZ was calculated based on their body weight. Before PTZ administration, mice were pre-injected IP with either the compounds [GAT1508 (60 mg/kg), GiGA1(60 mg/kg), ML297(60 mg/kg), valproic acid (VPA; 200 mg/kg) or CBD (at the concentrations indicated)] or their respective vehicles. Following a 30-minute video recording of the baseline behavior, the mice received a IP injection of PTZ at 60 mg/kg (Fig. 1). Video was recorded again for 30 min after PTZ injection, and the seizures were classified according to the modified Racine score (Table S1).

#### Drug formulation

GAT1508, ML297 and valproic acid (VPA) were prepared at 8 mg/ml concentration in vehicle consisting of: 5% DMSO, 10% Solutol, 85% of 10% Kleptose in Normal Saline (NS) GiGA1 was dissolved at 3 mg/ml concentration in vehicle consisting of: 10% DMSO, 40% PEG400, 5% Solutol and 45% Deionized (DI) Water. CBD was dissolved in a vehicle consisting of DMSO Tween 80 and normal saline in a ratio of 1:1:18 (total 20 parts).

Preparation steps: 1) Drug was first dissolved in DMSO and vortexed for 2 mins, 2) This was followed by 5 mins of Sonication, and 3) Each additional component was added sequentially with 2 mins of Vortexing.

Respective vehicle solution was used as a control to compare the results from the cohort injected with GAT1508, ML297, GiGA1 and VPA.

#### Molecular biology

Human GIRK1, and the highly homologous to human (96.2% identical) mouse GIRK2a were subcloned in pXOOM (24) while GABABR1a, GABABR2 were subcloned in pGEMHJ vectors. A single-cut restriction enzyme targeting the region following the b-globin 3’UTR of the plasmid was used to linearize the circular DNA. Linearized DNA was subjected to in vitro transcription using mMESSAGE mMACHINE T7 transcription kit (Thermo Fisher Scientific). Mutations were introduced by designing desired single point mutation primers. Mutagenesis was performed using the QuikChange site directed mutagenesis protocol and confirmed by restriction enzyme analysis. All the mutations were verified by sequencing.

#### *Xenopus laevis* oocyte expression

Survival surgeries were conducted on *Xenopus laevis* frogs to extract the oocytes. All procedures performed were approved by the Institutional Animal Care and Use Committee (IACUC) of Northeastern University, Boston, MA. The extracted oocytes were dissociated and defolliculated by collagenase treatment and for the experiments Stage V or VI oocytes were selected, and 2 ng of RNA was injected using a Nanoject Injector (Drummond Scientific). A final volume of 2.5 mL mixture containing RNA and DEPC treated water was made and filled into the glass injection needle (7-inch Drummond #3-000-203-G/XL). The needle was pulled using a P-97 micropipette puller (Sutter Instruments) to achieve a tip diameter of about 10-15 mM wide. 50 nL of the RNA mixture is injected into each oocyte. Injected oocytes were incubated at 17 °C for 36 h in OR2(++) solution. The standard OR2(++) solution contained 85 mM NaCl, 50 mM HEPES, 50 mM KCl, 50 mM NaOH, 10 mM MgCl_2_.6H_2_O. After an incubation period of 24-48 hours the oocytes were tested using TEVC.

#### Two-electrode voltage-clamp (TEVC)

Whole-cell currents in oocytes were measured by TEVC using a GeneClamp 500 (Molecular Devices) amplifier (25). The electrodes used for TEVC were pulled and filled with 3 M KCl in 1.5% (w/v0 agarose). The resistance of the electrodes was maintained between 0.5-1 MΩ. Potassium (K^+^) currents were measured in Low-K (NaCl 96 mM, HEPES 50 mM, KCl 20 mM, CaCl_2_.2H_2_O 18 mM, MgCl_2_.6H_2_O 10 mM) high-K^+^ (NaCl 20 mM, HEPES 50 mM, KCl 96 mM, CaCl_2_.2H_2_O 18 mM, MgCl_2_.6H_2_O 10 mM) and Drug (diluted in high-K^+^). Oocytes were held at 0 mV while a ramp protocol from −80 mV to +80 mV was applied every second. Currents were determined at -80 mV. Win WCP software (University of Strathclyde) was used to record the currents. 8-10 oocytes were studied for each condition.

#### Cryo-EM data collection and processing

GIRK1/2 was expressed in insect Sf9 cells and purified as described previously (16). GIRK1/2 proteins were prepared for cryo-EM at a concentration of 4.5 mg/ml with 0.5 mM diC8-PIP_2_ and 50 mM GAT1508.

Samples were first evaluated by negative-stain EM prior to cryo-EM grid preparation (26). Concentrated protein samples were applied to R1.2/1.3 UltrAuFoil grids and vitrified using a Vitrobot Mark IV (Thermo Fisher Scientific). Cryo-EM data were collected on a 300 kV Titan Krios transmission electron microscope (Thermo Fisher Scientific) equipped with BioQuantum energy filter and a Gatan K3 Direct Electron Detector, with automated acquisition controlled by EPU software. Each movie was recorded as 50 frames with a total dose of 50 e-/Å2 at a calibrated pixel size of 0.92 Å and a defocus range of -0.8 to -1.8 µm.

A total of 8,061 movie micrographs were collected for 3D reconstruction. After multi-rounds of particle picking, 2D classification, and 3D classification in RELION (27) and CryoSPARC (28), 48,276 particles were retained for final reconstruction. This yielded a 2.7 Å map with well-resolved density for GAT1508 inserted into the transmembrane domain (Table S2).

#### Model building and refinement

AlphaFold-predicted models of GIRK1 and GIRK2 were retrieved from the UniProt database and fitted into the final cryo-EM density map using UCSF ChimeraX (29). The models were manually rebuilt in Coot (30) and refined in real space with Phenix (31). Final model quality was assessed with MolProbity (32).

#### Brain slice electrophysiology

C57BL/6J mice (P30) were decapitated under Isoflurane anesthesia. All procedures performed were approved by the Institutional Animal Care and Use Committee (IACUC) of Northeastern University, Boston, MA. The brain was promptly removed and placed in ice-cold oxygenated aCSF containing (in mM): 124 NaCl, 3 KCl, 2 CaCl2, 2 MgSO4, 1.25 NaH2PO4, 26 NaHCO3, and 10 glucose, pH was adjusted to 7.4 using Hydrochloric acid (HCL). Horizontal slices (300 μm) of the hippocampal brain region were obtained using a vibratome, the slices were then allowed to equilibrate for 1 h in aCSF before recording.

The number of spikes in each voltage was analyzed using IGOR64 software to calculate the percent of spontaneous events.

#### Bliss-independence synergy analysis

To test whether the two compounds interact supra-additively, we compared the observed inhibition of the co-application against the Bliss-independence prediction computed from the effect of each compound separately. Spontaneous activity under perfusion of each drug was normalized to the mean High K^+^ response to yield a fractional inhibition, E = 1 − (mean_drug / mean_High K^+^). The expected inhibition of the combination under Bliss independence was computed as shown in the equation:

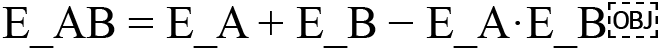

and the interaction term was defined as the observed minus the predicted combination inhibition (33). Confidence in the interaction was obtained by nonparametric bootstrap (34). Each group was resampled with replacement at its original size and the interaction recomputed over 100000 iterations, from which 95% confidence intervals (percentile and bias-corrected and accelerated, BCa) and a two-sided p-value was derived.

#### Gating molecular dynamics simulations

Molecular dynamics simulations were carried out on wild-type GIRK1/2 (WT-G1G2) and GAT1508 bound to GIRK1/2 (GAT-G1G2) systems in Amber22 using ff19SB forcefield for proteins and Lipids21 (35) parameters for lipids. Potassium chloride (150 mM) as well as neutralizing counterions were added to the systems. The CHARMM-GUI membrane builder web server was used to generate POPC:POPE:POPS:cholesterol (25:5:5:1) lipid bilayer for the systems. A TIP3P water box was used to solvate the complexes. The system was built using the tleap module in Amber, followed by hydrogen-mass repartitioning (HMR) to enable a 4-fs integration timestep during production run. Long-range electrostatics were treated with the Particle-mesh Ewald (PME) and periodic boundary conditions. Energy minimization was performed by keeping the solute restrained with a harmonic force constant of 500 kcal mol^−1^ Å^−2^ for 100 steps followed by 2000 steps of steepest descent. The system was heated gradually from 0 K to 100 K over 2500 steps using Langevin thermostat under constant volume while keeping proteins and lipids weakly restrained (500 kcal mol^−1^ Å^−2^ for the solute; 10 kcal mol^−1^ Å^−2^ for lipids). Subsequently, the temperature was gradually increased from 100 K to 303 K over 50,000 steps using anisotropic pressure coupling to reach 1 atm. The systems were equilibrated at 303 K under NPT conditions using Berendsen barostat and Langevin thermostat. SHAKE constraints were applied to all bonds and a time step of 1 fs was used. This was followed by a series of ten consecutive equilibration runs of 500 ps each (5 ns total) conducted at 303 K using a 2-fs time step. Weak restraints (10-20 kcal·mol^−1^·Å^−2^) were applied to protein backbone to preserve stable membrane-protein packing during equilibration. Each of these equilibrated systems WT-G1/2 and GAT-G1/2 were each subjected to 10 μs production runs, respectively, under NVT conditions. A static electric field with the magnitude of 0.138 kcal·mol^−1^·Å^−1^ (0.06 V/nm, 200 mV across membrane) was applied along the negative z-direction. The results of the simulations were analyzed using various tools, including ChimeraX (29), VMD (36) as well as by in-house scripts.

#### Interaction pattern analysis

CPPTRAJ and custom Python-based analysis scripts (written in pandas, numpy, and matplotlib) were utilized to explore key ligand-residue interactions throughout the simulation runs. Pairwise hydrogen-bond distances were calculated between critical protein residues and ligand donor/acceptor atoms. These included interactions involving Glu (E141.1), Asp (D173.1), Trp (W95.1) and Ser residues in GIRK2 (S177.2 or S181.2). The hydrogen-bond distances were monitored between ligand amide NH groups and carboxylate oxygens of Asp and Glu, the ligand oxygen of carbonyl with Trp, and the ligand nitrogen of the pyrazole ring with either Ser of GIRK2. The contact was considered as hydrogen-bond when the donor-acceptor distance was ≤ 3.5 Å. Permeation events were also recorded and mapped with the interaction pattern data to compute total simulation time and frequency of ion permeations associated with each ligand interaction pattern.

## Author contributions

S.M.S., J.S., T.K., D.E.L., K.S., G.S., L.D. P., and M.C. designed the study. J.S. led the execution of all in vivo work with N.C-R and L.V. and led the voltage clamp studies. A.Z. performed voltage-clamp and mutagenesis experiments. L.W. performed cryo-EM sample preparation, data collection, processing, model-building and structural analysis together with G.S. T.K. purified proteins and prepared all mutants. L.V and N.R. performed hippocampal slice recordings. A.S.-C., and E.D. carried out patch-clamp experiments and K.W.C and L.C. performed TEVC recordings. N.P. and M.Z. performed gating molecular dynamics simulations and calculations. R.M., E.D., L.C., W.F., and G.A.T. supervised or synthesized the urea compounds used in the study. S.M.S. and D.E.L. conceived and supervised the study. J.S., L.W. M.C. and S.M.S. prepared figures. J.S., D.E.L., and S.M.S. wrote the manuscript with input from all authors.

## Acknowledgments

This work was supported by NIH R01 grants HL059949-28 and NS131467-03 to D.E.L and DoD CDMRP ERP EP230110 to S.M.S. We thank Heikki Vaananen for electrophysiology technical guidance and administrative support; Mahmoud Nasr (Brigham and Women’s Hospital/Harvard Medical School), Sami Noujaim (University of South Florida), and Satya Kunapuli (Temple University); and members of the Logothetis and Plant laboratories for discussions. Computational resources were provided by Research Computing within Information Technology Services at Northeastern University and by the Argonne Leadership Computing Facility at Argonne National Laboratory.

---

**Table S1.** Modified Racine Scores.

| Score | Behavioral Expression |
| --- | --- |
| 0 | Normal; |
| 1 | immobilization/sudden behavioral arrest, sniffing; |
| 2 | head nodding, facial and forelimb clonus (short myoclonic jerk); |
| 3 | continuous myoclonic jerk, tail rigidity, stiffness; (clonic activity not generalized); |
| 4 | generalized limbic seizures with kangaroo posture or violent convulsions; (righting reflex present); |
| 5 | continuous generalized seizures, lying on belly or on side, wild jumping, righting reflex lost (tonic or clonic-tonic convulsions); |
| 6 | tonic extension or death |

**Table S2.**
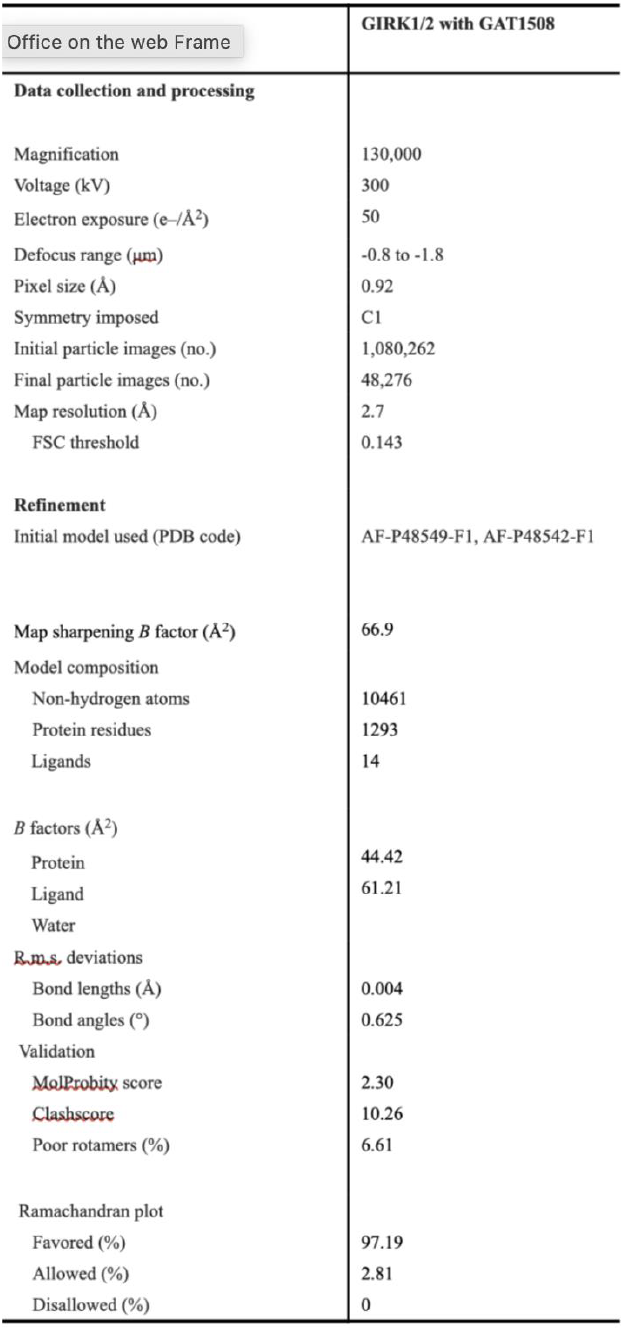
Croy-EM data collection, refinement and validation station statistics.

| Office on the web Frame | GIRK1/2 with GAT1508 |
| --- | --- |
| **Data collection and processing** |  |
| Magnification | 130,000 |
| Voltage (kV) | 300 |
| Electron exposure (e-/Å<sup>2</sup>) | 50 |
| Defocus range (μm) | -0.8 to -1.8 |
| Pixel size (Å) | 0.92 |
| Symmetry imposed | C1 |
| Initial particle images (no.) | 1,080,262 |
| Final particle images (no.) | 48,276 |
| Map resolution (Å) | 2.7 |
| FSC threshold | 0.143 |
| **Refinement** |  |
| Initial model used (PDB code) | AF-P48549-F1, AF-P48542-F1 |
| **Map sharpening *B* factor (Å<sup>2</sup>)** | 66.9 |
| Model composition |  |
| Non-hydrogen atoms | 10461 |
| Protein residues | 1293 |
| Ligands | 14 |
| *B* factors (Å<sup>2</sup>) |  |
| Protein | 44.42 |
| Ligand | 61.21 |
| Water |  |
| ~~R.m.s.~~ deviations |  |
| Bond lengths (Å) | 0.004 |
| Bond angles (°) | 0.625 |
| Validation |  |
| ~~Mol~~Probity score | 2.30 |
| ~~Clash~~score | 10.26 |
| Poor rotamers (%) | 6.61 |
| Ramachandran plot |  |
| Favored (%) | 97.19 |
| Allowed (%) | 2.81 |
| Disallowed (%) | 0 |

## References

1. Logothetis DE, Kurachi Y, Galper J, Neer EJ, Clapham DE. The beta gamma subunits of GTP-binding proteins activate the muscarinic K+ channel in heart. Nature. 1987 Jan 22-28;325(6102):321–6. doi: 10.1038/325321a0.

2. Reuveny E, Slesinger PA, Inglese J, Morales JM, Iñiguez-Lluhi JA, Lefkowitz RJ, Bourne HR, Jan YN, Jan LY. Activation of the cloned muscarinic potassium channel by G protein beta gamma subunits. Nature. 1994 Jul 14;370(6485):143–6. doi: 10.1038/370143a0.

3. Sui JL, Chan KW, Logothetis DE. Na+ activation of the muscarinic K+ channel by a G-protein-independent mechanism. J Gen Physiol. 1996 Nov;108(5):381–91. doi: 10.1085/jgp.108.5.381.

4. Sui JL, Petit-Jacques J, Logothetis DE. Activation of the atrial KACh channel by the betagamma subunits of G proteins or intracellular Na+ ions depends on the presence of phosphatidylinositol phosphates. Proc Natl Acad Sci U S A. 1998 Feb 3;95(3):1307–12. doi: 10.1073/pnas.95.3.1307.

5. Huang CL, Feng S, Hilgemann DW. Direct activation of inward rectifier potassium channels by PIP2 and its stabilization by Gbetagamma. Nature. 1998 Feb 19;391(6669):803–6. doi: 10.1038/35882.

6. Li, D., Jin, T., Gazgalis, D., Cui, M., Logothetis, D. E. (2019) Mechanism of GIRK2 gating by PIP2, Na+, and Gβγ. J. Biol. Chem. 294, 18934–18948.

7. Gazgalis D, Cantwell L, Xu Y, Thakur GA, Cui M, Guarnieri F, Logothetis DE. Use of a Molecular Switch Probe to Activate or Inhibit GIRK1 Heteromers In Silico Reveals a Novel Gating Mechanism. Int J Mol Sci. 2022 Sep 16;23(18):10820. doi: 10.3390/ijms231810820.

8. Cui, M., Cantwell, L., Zorn, A., Logothetis, D. E. (2021) Kir channel molecular physiology, pharmacology, and therapeutic implications. Handb. Exp. Pharmacol. 267, 277–356.

9. Lüscher, C., Slesinger, P. A. (2010) Emerging roles for GIRK channels in health and disease. Nat. Rev. Neurosci. 11, 301–315.

10. Luján, R., Marron Fernandez de Velasco, E., Aguado, C., Wickman, K. (2014) New insights into the therapeutic potential of GIRK channels. Trends Neurosci. 37, 20–29.

11. Petit-Jacques J, Sui JL, Logothetis DE. Synergistic activation of G protein-gated inwardly rectifying potassium channels by the betagamma subunits of G proteins and Na+ and Mg2+ ions. J Gen Physiol. 1999 Nov;114(5):673–84. doi: 10.1085/jgp.114.5.673.

12. Xu Y, Cantwell L, Molosh AI, Plant LD, Gazgalis D, Fitz SD, Dustrude ET, Yang Y, Kawano T, Garai S, Noujaim SF, Shekhar A, Logothetis DE, Thakur GA. The small molecule GAT1508 activates brain-specific GIRK1/2 channel heteromers and facilitates conditioned fear extinction in rodents. J Biol Chem. 2020 Mar 13;295(11):3614–3634. doi: 10.1074/jbc.RA119.011527. Epub 2020 Jan 17.

13. Kaufmann, K., Romaine, I., Days, E., et al. (2013) ML297 displays antiepileptic properties in mice. ACS Chem. Neurosci. 4, 1278–1286.

14. Zhao, Y., Ung, P. M., Zahoránszky-Kőhalmi, G., et al. (2020) Identification of a G-protein-independent activator of GIRK channels. Cell Rep. 31, 107770.

15. Wang, L., Kawano, T., Santa-Cruz, A., Parvaiz, N., Zorn, A., Lu, Y., Zaka, M., Aly, N., Pentikis, J., Dadiotis, E., et al. Stoichiometry-dependent gating and structural pharmacology of cardiac GIRK1/4 channels. Under review.

16. Kawano, T., Wang, L., Chan, K. W., Zorn, A., Parvaiz, N., Zaka, M., Santa-Cruz, A., Lu, Y., Aly, N., Pentikis, J., et al. Asymmetric assembly and activation mechanism of heterotetrameric GIRK1/2 channels. Under review.

17. Wang L, Wu Y, Deng Y, Kim B, Pierce L, Krilov G, Lupyan D, Robinson S, Dahlgren MK, Greenwood J, Romero DL, Masse C, Knight JL, Steinbrecher T, Beuming T, Damm W, Harder E, Sherman W, Brewer M, Wester R, Murcko M, Frye L, Farid R, Lin T, Mobley DL, Jorgensen WL, Berne BJ, Friesner RA, Abel R. Accurate and reliable prediction of relative ligand binding potency in prospective drug discovery by way of a modern free-energy calculation protocol and force field. J Am Chem Soc. 2015 Feb 25;137(7):2695–703. doi: 10.1021/ja512751q. Epub 2015 Feb 12.

18. Cui M, Lu Y, Logothetis DE. Understanding Ion Channel Gating Mechanisms by Molecular Dynamics Simulations. Adv Exp Med Biol. 2026;1497:19–31. doi: 10.1007/978-3-032-07523-9_2.

19. Li, D. L., Hu, L., Wang, L., Chen, C. L. (2020) Permeation mechanisms of GIRK2. Comput. Struct. Biotechnol. J. 18, 3950–3958.

20. Li, D., Shi, D., Wang, L. (2024) Structural insights into GIRK2 permeation. Biochim. Biophys. Acta Biomembr.1866, 184231.

21. Russo EB, Burnett A, Hall B, Parker KK. Agonistic properties of cannabidiol at 5-HT1a receptors. Neurochem Res. 2005 Aug;30(8):1037–43. doi: 10.1007/s11064-005-6978-1.

22. Jeong HJ, Han SH, Min BI, Cho YW. 5-HT1A receptor-mediated activation of G-protein-gated inwardly rectifying K+ current in rat periaqueductal gray neurons. Neuropharmacology. 2001 Aug;41(2):175–85. doi: 10.1016/s0028-3908(01)00062-4.

23. Devinsky O, Cross JH, Wright S. Trial of Cannabidiol for Drug-Resistant Seizures in the Dravet Syndrome. N Engl J Med. 2017 Aug 17;377(7):699–700. doi: 10.1056/NEJMc1708349. PMID: 28813226.

24. Jespersen T, Grunnet M, Angelo K, Klaerke DA, Olesen SP. Dual-function vector for protein expression in both mammalian cells and Xenopus laevis oocytes. Biotechniques. 2002 Mar;32(3):536–8, 540. doi: 10.2144/02323st05.

25. Hatcher-Solis C, Fribourg M, Spyridaki K, Younkin J, Ellaithy A, Xiang G, Liapakis G, Gonzalez-Maeso J, Zhang H, Cui M, Logothetis DE. G protein-coupled receptor signaling to Kir channels in Xenopus oocytes. Curr Pharm Biotechnol. 2014;15(10):987–95. doi: 10.2174/1389201015666141031111916.

26. Peisley, A., and Skiniotis, G. (2015). 2D Projection Analysis of GPCR Complexes by Negative Stain Electron Microscopy. Methods Mol Biol 1335, 29–38. 10.1007/978-1-4939-2914-6_3.

27. Kimanius, D., Dong, L., Sharov, G., Nakane, T., and Scheres, S.H.W. (2021). New tools for automated cryo-EM single-particle analysis in RELION-4.0. Biochem J 478, 4169–4185. 10.1042/BCJ20210708.

28. Punjani, A., Rubinstein, J.L., Fleet, D.J., and Brubaker, M.A. (2017). cryoSPARC: algorithms for rapid unsupervised cryo-EM structure determination. Nat Methods 14, 290–296. 10.1038/nmeth.416910.1038/nmeth.4169.

29. Meng, E.C., Goddard, T.D., Pettersen, E.F., Couch, G.S., Pearson, Z.J., Morris, J.H., and Ferrin, T.E. (2023). UCSF ChimeraX: Tools for structure building and analysis. Protein Sci 32, e4792. 10.1002/pro.4792.

30. Emsley, P., Lohkamp, B., Scott, W.G., and Cowtan, K. (2010). Features and development of Coot. Acta Crystallogr D Biol Crystallogr 66, 486–501. 10.1107/S0907444910007493.

31. Liebschner, D., Afonine, P.V., Baker, M.L., Bunkóczi, G., Chen, V.B., Croll, T.I., Hintze, B., Hung, L.W., Jain, S., McCoy, A.J., et al. (2019). Macromolecular structure determination using X-rays, neutrons and electrons: recent developments in Phenix. Acta Crystallogr D Struct Biol 75, 861–877. 10.1107/S2059798319011471.

32. Chen, V.B., Arendall, W.B. 3rd, Headd, J.J., Keedy, D.A., Immormino, R.M., Kapral, G.J., Murray, L.W., Richardson, J.S., and Richardson, D.C. (2010). MolProbity: all-atom structure validation for macromolecular crystallography. Acta Crystallogr D Biol Crystallogr 66, 12–21. 10.1107/S0907444909042073

33. Demidenko E, Miller TW. Statistical determination of synergy based on Bliss definition of drugs independence. PLoS One. 2019 Nov 25;14(11):e0224137. doi: 10.1371/journal.pone.0224137. PMID: 31765385; PMCID: PMC6876842.

34. Van der Borght K, Tourny A, Bagdziunas R, Thas O, Nazarov M, Turner H, Verbist B, Ceulemans H. BIGL: Biochemically Intuitive Generalized Loewe null model for prediction of the expected combined effect compatible with partial agonism and antagonism. Sci Rep. 2017 Dec 20;7(1):17935. doi: 10.1038/s41598-017-18068-5. PMID: 29263342; PMCID: PMC5738392.

35. Dickson, C. J., Madej, B. D., Skjevik, A. A., Betz, R. M., Teigen, K., Gould, I. R.,, and Walker, R. C. (2022). Lipid21: Complex Lipid Membrane Simulations with AMBER. Journal of Chemical Theory and Computation, 18(3), 2035–2048

36. Humphrey, W., Dalke, A., and Schulten, K. (1996). VMD – Visual molecular dynamics. Journal of Molecular Graphics, 14(1), 33–38.

