## Supplementary Figures and Legends for "GAT1508, a selective neuronal GIRK1/2 activator, exhibits superior antiepileptic efficacy and synergizes with cannabidiol"

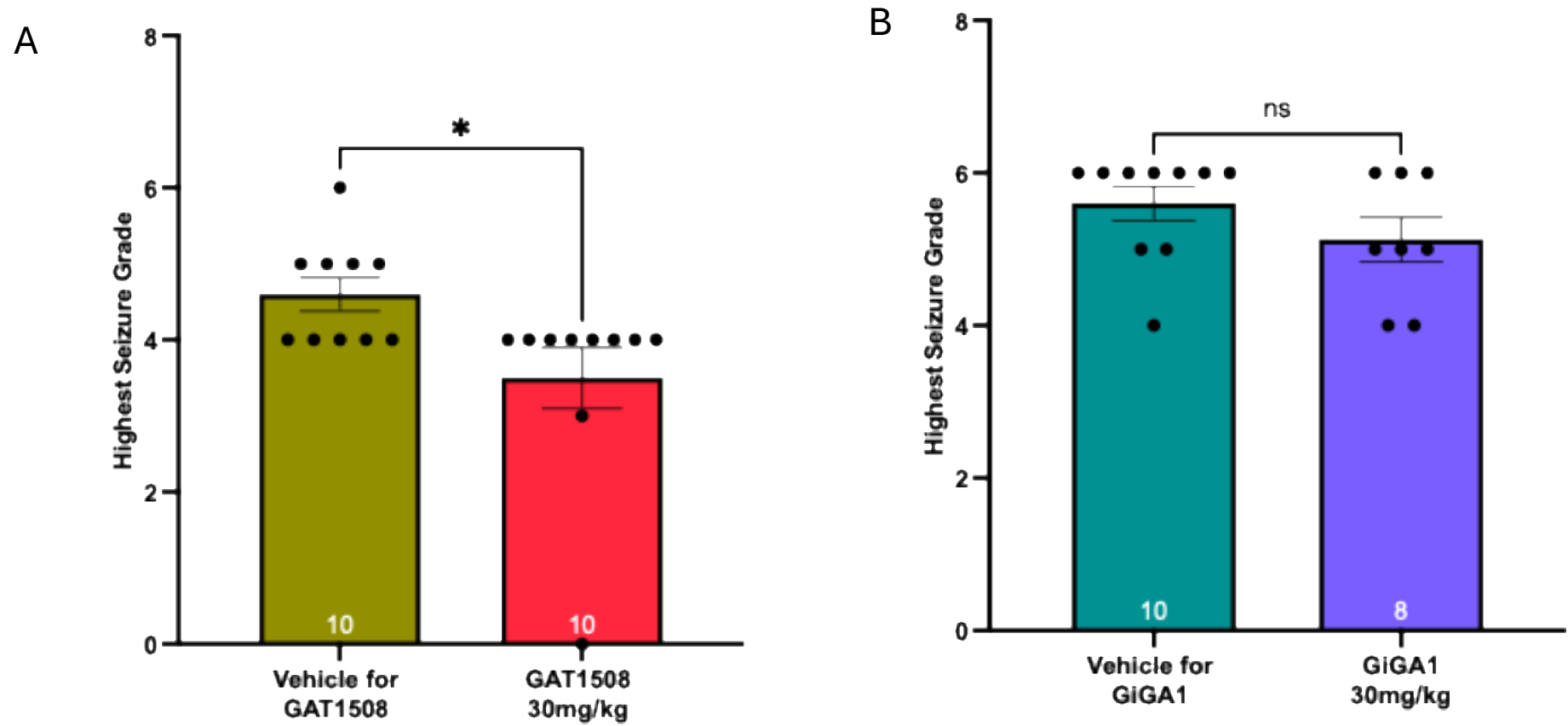

**Figure S1 : GAT1508 at 30mg/kg was significantly effective compared to GiGA1 at 30mg/kg.**

PTZ induced seizure experiments were performed in C57BL/6 mice using half the doses of GAT1508 and GiGA1 that were evaluated in previous experiments. Different formulations were used for both the compounds, and each compound was compared to its respective vehicle control. *A*, GAT1508 (30mg/kg) markedly reduced seizure severity, with no animals exhibiting any seizures above grade 4, demonstrating robust antiseizure efficacy even at 30mg/kg. *B*, In contrast, GiGA1 administered at the corresponding dose did not significantly reduce seizure compared with its vehicle control.

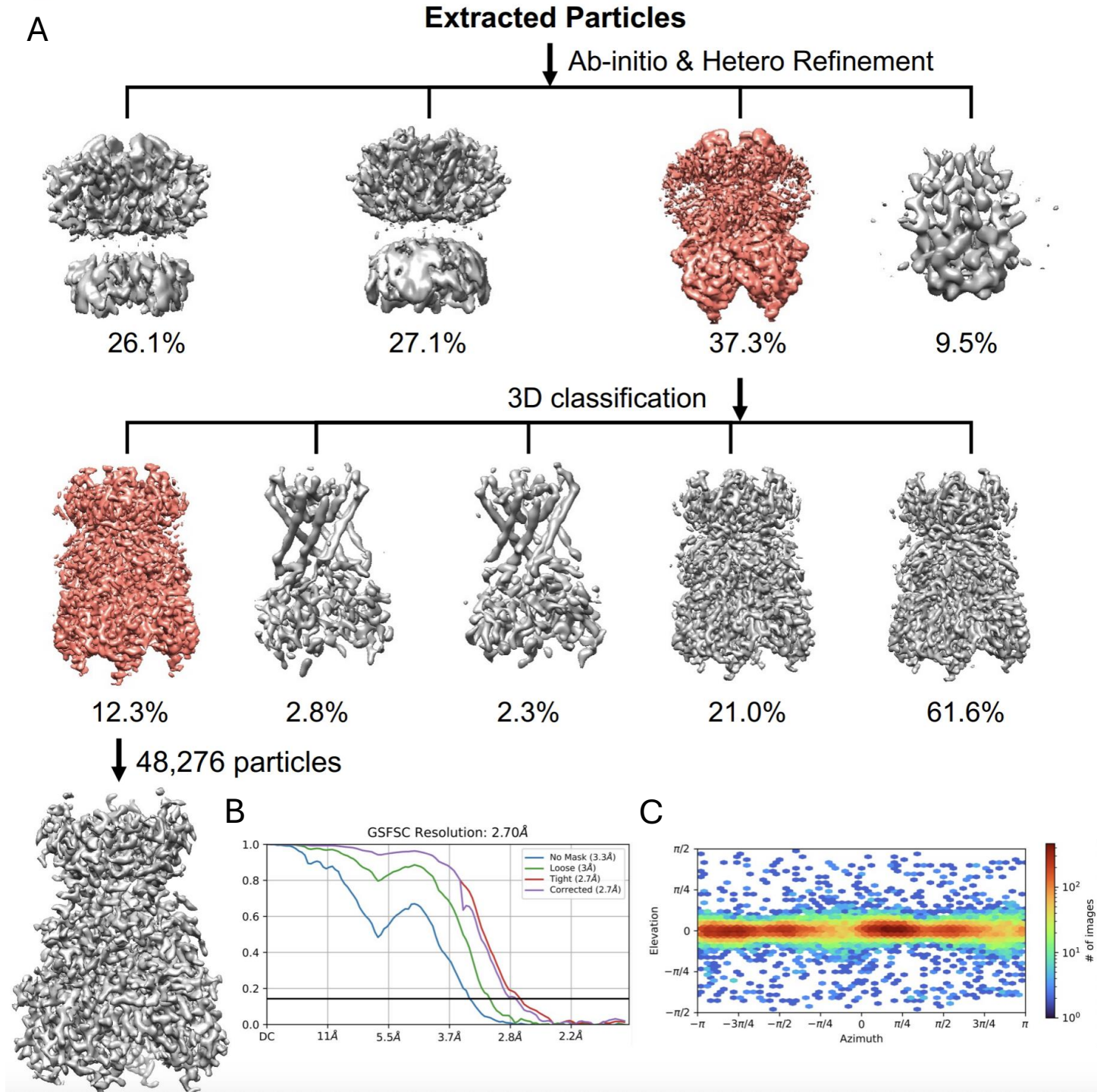

Fig. S2 Cryo-EM processing and reconstruction of GAT1508-bound GIRK1/2 complex.

**A**, Summary of image processing procedures for GIRK1/2 with compound GAT1508.

**B**, Fourier shell correlation (FSC) curves between the two half-maps.

**C**, Angular distribution plots for the final reconstruction.

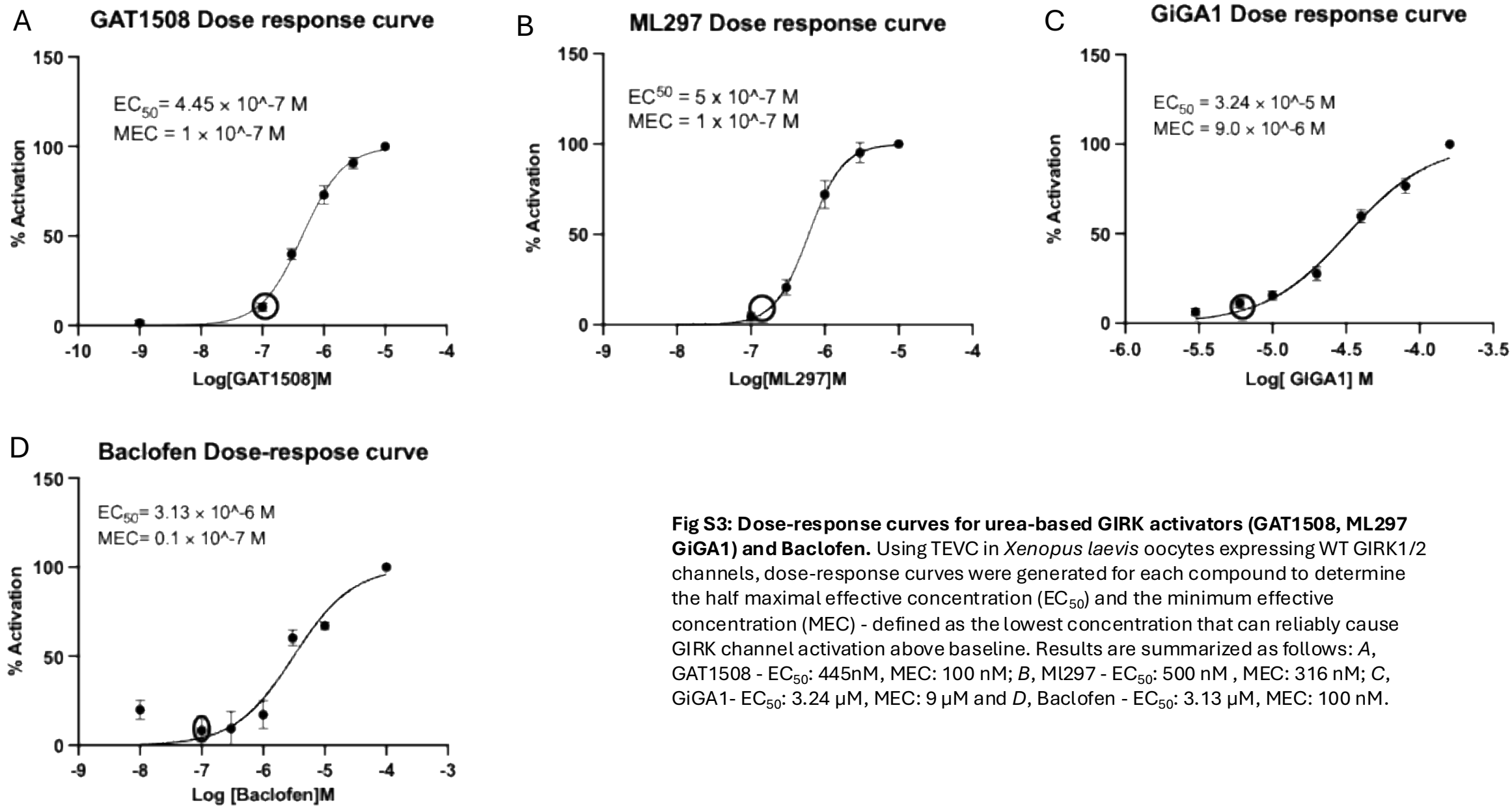

**Fig S3: Dose-response curves for urea-based GIRK activators (GAT1508, ML297 GiGA1) and Baclofen.** Using TEVC in *Xenopus laevis* oocytes expressing WT GIRK1/2 channels, dose-response curves were generated for each compound to determine the half maximal effective concentration (EC<sub>50</sub>) and the minimum effective concentration (MEC) - defined as the lowest concentration that can reliably cause GIRK channel activation above baseline. Results are summarized as follows: A, GAT1508 - EC<sub>50</sub>: 445nM, MEC: 100 nM; B, ML297 - EC<sub>50</sub>: 500 nM, MEC: 316 nM; C, GiGA1- EC<sub>50</sub>: 3.24 μM, MEC: 9 μM and D, Baclofen - EC<sub>50</sub>: 3.13 μM, MEC: 100 nM.
